# The neural dynamics of familiar face recognition

**DOI:** 10.1101/393652

**Authors:** Géza Gergely Ambrus, Daniel Kaiser, Radoslaw Martin Cichy, Gyula Kovács

**Author notes:** These authors contributed equally to the study (Géza Gergely Ambrus, Daniel Kaiser).

## Abstract

In real-life situations, the appearance of a person’s face can vary substantially across different encounters, making face recognition a challenging task for the visual system. Recent fMRI decoding studies have suggested that face recognition is supported by identity representations located in regions of the occipito-temporal cortex. Here, we used EEG to elucidate the temporal emergence of these representations. Human participants (both sexes) viewed a set of highly variable face images of four highly familiar celebrities (two male, two female), while performing an orthogonal task. Univariate analyses of event-related EEG responses revealed a pronounced differentiation between male and female faces, but not between identities of the same sex. Using multivariate representational similarity analysis, we observed a gradual emergence of face identity representations, with an increasing degree of invariance. Face identity information emerged rapidly, starting shortly after 100ms from stimulus onset. From 400ms after onset and predominantly in the right hemisphere, identity representations showed two invariance properties: (1) they equally discriminated identities of opposite sexes and of the same sex, and (2) they were tolerant to image-based variations. These invariant representations may be a crucial prerequisite for successful face recognition in everyday situations, where the appearance of a familiar person can vary drastically.

**Significance Statement:** Recognizing the face of a friend on the street is a task we effortlessly perform in our everyday lives. However, the necessary visual processing underlying familiar face recognition is highly complex. As the appearance of a given person varies drastically between encounters, for example across viewpoints or emotional expressions, the brain needs to extract identity information that is invariant to such changes. Using multivariate analyses of EEG data, we characterize how invariant representations of face identity emerge gradually over time. After 400ms of processing, cortical representations reliably differentiated two similar identities (e.g., two famous male actors), even across a set of highly variable images. These representations may support face recognition under challenging real-life conditions.

## Introduction

Efficient face recognition is a key ability in human’s everyday lives, and many studies have investigated its underlying neural mechanisms (Gobbini and Haxby, 2007; Duchaine and Yovel, 2015). Recently, much progress has been made in spatially pinpointing the neural correlates of face recognition by advances in multivariate classification techniques for fMRI data (Anzellotti and Caramazza, 2014). These techniques have allowed researchers to decode face identity from different regions of the face processing network, such as from the fusiform face area (FFA; Gilaie-Dotan and Malach, 2007; Nestor et al., 2011; Goesaert and Op de Beeck, 2013; Verosky et al., 2013; Anzellotti et al., 2014; Axelrod and Yovel, 2015; Weibert et al., 2016), the anterior temporal lobe (ATL; Kriegeskorte et al., 2007; Nasr and Tootell, 2012; Anzellotti et al., 2014) or from a larger network extending from early visual areas towards the inferior frontal gyrus (Visconti Di Oleggio Castello et al., 2017).

The temporal emergence of face identity representations, however, remains relatively unexplored. Most of our knowledge on the temporal dynamics of face recognition stems from EEG and magnetoencephalography (MEG) studies employing traditional, univariate analyses on temporally confined ERP/MEP components. Across these studies, the components associated with face recognition vary substantially: Several reports have linked face recognition to the P100 and N170 components (Debruille et al., 1998; Heisz et al., 2006; Caharel et al., 2009; Rousselet et al., 2009; Liu et al., 2013), others have stressed the role of the later N250 and N400 components (Bentin and Deouell, 2000; Schweinberger et al., 2002; Huddy et al., 2003; Tanaka et al., 2006; Curran and Hancock, 2007; Gosling and Eimer, 2011; Jin et al., 2012).

So far only three studies have used multivariate pattern analysis (MVPA) to evaluate the temporal dynamics of face identity processing (Nemrodov et al., 2016, 2018; Vida et al., 2017). All three investigated the temporal emergence of identity representations across changes in emotional expression, revealing that identity representations emerge relatively early within the first 200ms after stimulus onset.

However, these previous studies suffer from two critical shortcomings. First, they used unfamiliar faces, whose processing is assumed to be markedly different from the processing of familiar faces, as reflected both in behavioral performance (Johnston and Edmonds, 2009) and neural activations (Natu and O’Toole, 2011). Second, variability across images of the same identity was very limited, leaving it unclear how their results generalize to everyday face recognition where individual encounters with highly-variable, “ambient” face images give rise to drastic visual differences (Mike Burton, 2013; Young and Burton, 2017; Kramer et al., 2018).

In the current EEG study, we provide a temporal characterization of face identity processing, which eliminates both shortcomings: First, we used images of four celebrities, who were highly familiar to the participants (Fig. 1A). Second, for each identity, we used 10 “ambient” images (Jenkins et al., 2011), which varied substantially in a range of properties, such as viewpoint, lighting, and expression.

**Figure 1.**
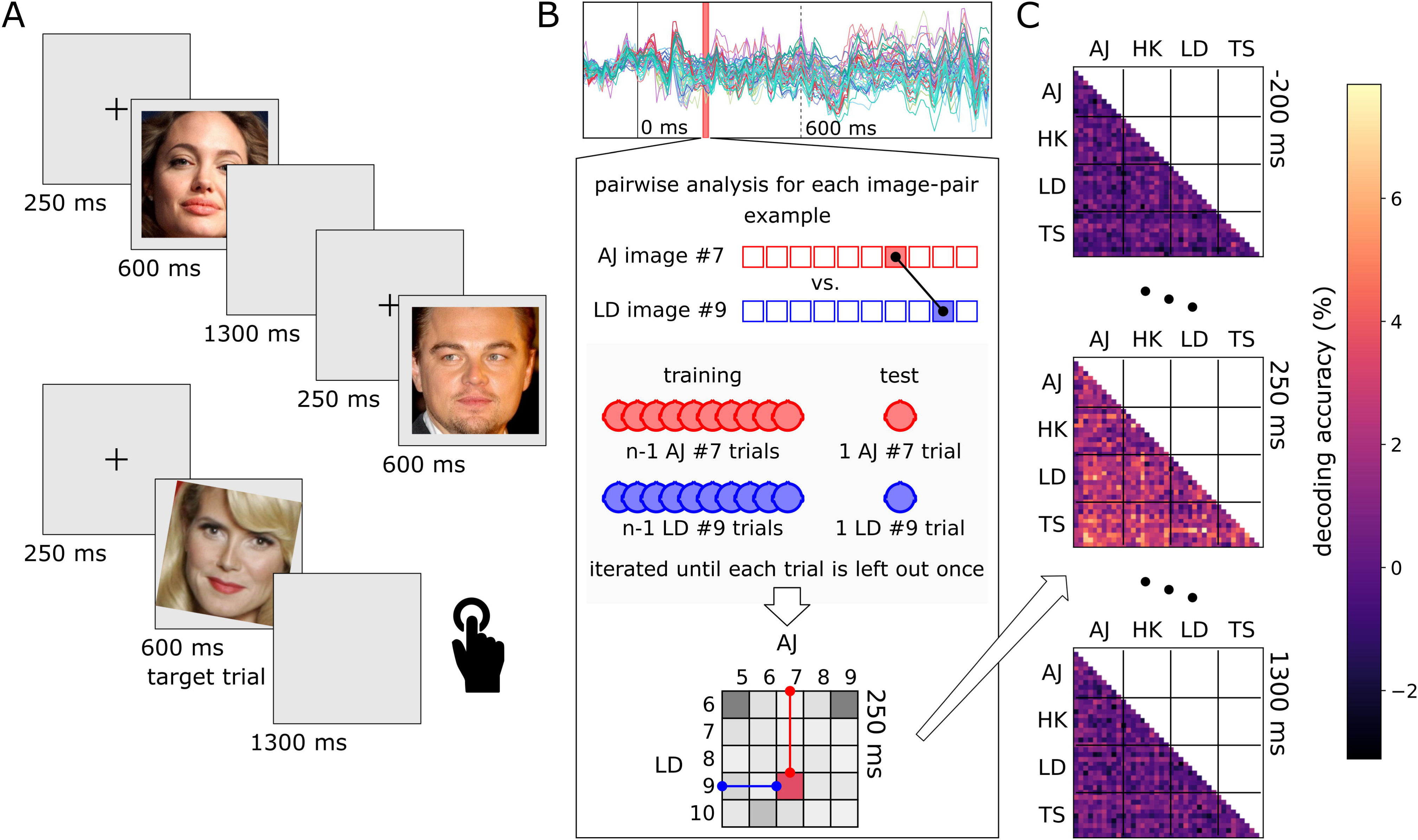
Design and Analysis Approach. A. Trial structure and stimulus examples. Stimuli were “ambient”, face-cropped images of four highly recognized celebrities (Angelina Jolie (AJ), Heidi Klum (HK), Leonardo DiCaprio (LD), Til Schweiger (TS))^1^. Each trial started with a fixation cross (250ms), followed by the stimulus image (600ms) and a blank screen (1300ms). Target trials containing a tilted stimulus (illustrated on the bottom) were included to ensure that participants maintained attention. B. The logic of the multivariate pattern analysis. Top: A representative ERP recording from one participant. EEGs were segmented between ‒200 to 1300ms relative to stimulus onset. Bottom: For each time point separately, linear classification analyses were performed for each combination of individual images, using a leave-one-trial-out scheme. This procedure resulted in a 40×40 matrix (i.e., 10 images for each of the 4 identities) of decoding accuracies at each time point. C. Representational dissimilarity matrices (RDMs) showing pairwise decoding accuracies at ‒200, 250 and 1300ms relative to stimulus onset.

Using representational similarity analysis (RSA; Kriegeskorte and Kievit, 2013), we show that the earliest representations of facial identity emerge shortly after 100ms post-stimulus and most robustly in posterior electrodes. Later representations, emerging from 400ms onwards and in electrodes over right occipito-temporal cortex, contained identity information for faces of the same sex and were invariant to image-based properties. Our results suggest that familiar face recognition is supported by fine-grained neural representations in the face processing network, where identity information over time becomes increasingly invariant to other visual and conceptual properties of the face.

## Methods

### Participants

Twenty-six healthy participants (6 male), with an average age of 25 years (SD = 5.0) took part in the study in exchange for partial course credits or monetary compensation. The experiment was conducted in accordance with the guidelines of the Declaration of Helsinki, and with the approval of the ethics committee of the University of Jena. Written informed consent was acquired from all participants.

### Stimuli

The stimuli were ambient, color photographs two female (Angelina Jolie, AJ; Heidi Klum, HK) and two male (Leonardo DiCaprio, LD; Til Schweiger, TS) celebrities. We selected these celebrities based on a pilot survey where we collected familiarity ratings across a range of well-known celebrities in Germany. For each identity, ten images were selected from a pool of web-scraped photographs, pre-screened for quality. Stimulus images were cropped to a rectangle centered on the inner features of the face (see Fig. 1A). To ensure substantial variation across images depicting the same identity, we selected images that minimized the structural similarity index (SSIM; Wang et al., 2004) among images of the same identity, while maximizing it among images of identities of the same sex; this was achieved by using random combination sorting with 100,000 iterations per sex category. The resulting mean SSIM values were: LD: 0.387, TS: 0.378, LD vs TS: 0.355; AJ: 0.379, HK: 0.371, AJ vs HK: 0.337. Stimuli were presented centrally on a uniform gray background on a TFT display (1680 × 1050 pixel resolution, refresh rate 60 Hz). The experiment was written in Psychopy (Peirce, 2008).

### Experimental procedure

A total of 1760 trials (1600 non-target and 160 target) were presented in 40 runs, each containing the ten images of the four identities once, in a pseudo-random order (with the constraint that the same identity was never repeated in two consecutive trials). Thus, photos of one identity were seen 400 times, so that a given image of a given identity was presented 40 times during the experiment. Each trial started with a fixation cross (250ms), followed by the stimulus image (600ms, subtending a visual angle of 4.4° in diameter) and finally a blank display (1300ms). Short breaks were provided after every 10 runs, but the run boundaries were not indicated otherwise. There were four target-trials in each run, where the image was rotated 10 degrees clockwise or anticlockwise. Participants were instructed to press the space bar when they saw a target image (the overall detection accuracy was at 99.52 ± 0.67%). These target trials served to ensure that the participants maintained their attention, and were not included in the analysis. An average experimental session lasted 81.5 (±5.3) minutes.

At the beginning of each experiment, prior to mounting the electrode caps, participants were presented images of the four identities and were asked to name them. All our participants were able to name all four celebrities correctly. The images of this initial familiarity-testing phase were not part of the later EEG experiment. After the EEG recording, participants were asked to rate their familiarity with the identities on a 7-point scale. Mean ratings were generally high (AJ: 6.12, HK: 6.30, LD: 6.15, TS: 6.11) and not statistically different for the four identities (F[3,75] = 0.243, p = 0.866).

### EEG recording and preprocessing

The EEG was recorded in a dimly lit, electrically shielded, and sound–attenuated chamber. The distance between the eyes and the computer screen was set to 96 cm via a chin rest. The electroencephalogram (EEG) was recorded with a 512 Hz sampling rate (bandwidth: DC to 120 Hz) using a 64-channel Biosemi Active II system. Electrooculogram (EOG) was recorded from the outer canthi of the eyes and from above and below the left eye.

The preprocessing pipeline was implemented in MNE-python (Gramfort et al., 2013, 2014). EEG was notch-filtered at 50 Hz, band-pass filtered between 0.1 and 70 Hz, segmented from ‒200 to 1300ms relative to stimulus onset, and baseline corrected with respect to the first 200ms. Artefact rejection was carried out using the “Autoreject” algorithm (Jas et al., 2017). The resulting data was downsampled to 100Hz to increase signal-to-noise ratio in the multivariate analyses (Grootswagers et al., 2017).

### Event-related potentials

To test for the presence of identity-related information within the conventional ERPs we averaged data across repetitions for each facial identity, electrode and participant separately. Next, we created grand-averages of these data across six regions of interests, corresponding to the left and right anterior (Fp, AF, F, FC), central (FT, TP, C, T) and posterior occipito-temporal electrodes (PO, P, O, I). The central electrodes (Fpz, AFz, Fz, FCz, Cz, CPz, Pz, POz, Oz, Iz) were included in both the left and the right clusters; this was done to maintain sufficient electrode counts for the multivariate analyses (see below). For reasons of consistency, the same electrode clusters were used in both analyses. The posterior clusters included the electrodes typically yielding the largest face-sensitive N170 components (Rossion and Jacques, 2008). First, we tested for identity selectivity by using a one-way repeated measures ANOVA with identity (4) as a factor. Second, we averaged the two female and two male face elicited ERPs and performed a paired t-test for testing sex-specific differences. Third, we tested if the ERPs differed for the two identities within the same sex by comparing the ERPs for the two female as well as for the two male identities with each other in t-tests.

### Representational Similarity Analysis

To model the neural organization of face representations, we performed a representational similarity analysis (RSA; Kriegeskorte et al., 2008) on the EEG data. In this analysis (Fig. 1B/C), the neural dissimilarity between all pairs of face images (i.e., between all 40 individual images), was modeled as a function of different predictor matrices (see below).

#### Neural dissimilarity

Neural dissimilarity was extracted by performing a linear classification analysis, where pairwise decoding accuracies were used as a measure of representational dissimilarity. Classification analysis was carried out using the CoSMoMVPA toolbox (Oosterhof et al., 2016). Linear-discriminant-analysis (LDA) classifiers were trained and tested on response patterns across all 64 electrodes, separately for each time point across the epoch (downsampled to 100 Hz, i.e., with a 10ms resolution) and separately for each pair of images. Training and testing was done in a leave-one-out scheme (Fig. 1B): classifiers were trained on all but one trials for each of the two conditions, and tested on the left-out trials. This procedure was repeated until each trial was left out once, and classification accuracy was averaged across these repetitions. Pairwise classification time-courses were smoothed with a 30ms (i.e., 3 consecutive time points) averaging window (Kaiser et al., 2016). This classification analysis led to one representational dissimilarity matrix (RDM; 40×40 entries, with empty diagonal) for each time point (Fig. 1C).

#### Modelling neural dissimilarity

To model the neural dissimilarity, we created four categorical predictor RDMs. Each predictor RDM covered 40×40 elements, and contained zeros where the entries represented comparisons of similar images (i.e., similar on the dimension of interest, see below) and ones, where the entries reflected comparisons of dissimilar images. To quantify correspondence between the predictor RDMs and the neural RDMs, we unfolded the lower off-diagonal elements of the matrices into two vectors (i.e., the diagonal of both matrices was discarded) and correlated the vectors using Spearman’s correlation coefficients. These correlations were computed separately for each time point, leading to a time series of correlations that reflected the correspondence of the neural data and the predictor. Individual-participant correlations were Fisher-transformed.

#### Modelling identity Information

For assessing differences between the four identities, all comparisons within a given identity (e.g., two images of AJ) were marked as similar (0) and all comparisons between two identities (e.g., an image of AJ and an image of TS) were marked as dissimilar (1) (Fig. 3A).

#### Modelling sex information

For assessing differences between face sexes, all comparisons within the same sex (e.g., an image of AJ and an image of HK) were marked as similar (0), and all comparisons between the different sexes (e.g., an image of AJ and an image of TS) were marked as dissimilar (1). To avoid confounding sex information with identity information, all comparisons within the same identity (e.g., two different images of AJ) were excluded from this analysis (as including these comparisons would overestimate the effect of sex) (Fig. 3C).

#### Modelling identity information between and within sexes

To uncover interactions between sex and identity processing, we constructed identity predictor RDMs that only covered all comparisons across the sexes or within one sex. The between-sex RDM was generated from the identity predictor matrix by removing all comparisons of two different identities of the same sex (e.g., an image of AJ and an image of HK), leaving only comparisons within identity (0) and between identities of the opposite sex (1). The within-sex RDM was generated from the identity predictor matrix by removing all comparisons of two identities of different sexes (e.g., an image of AJ and an image of TS), leaving only comparisons within identity (0) and between identities of the same sex (1) (Fig. 3E). Note that this within-sex analysis tests for identity representations in more thorough way: by removing between-sex comparisons, the more pronounced differences between faces of the opposite sex (due to face sex, and due to visual differences) are eliminated.

#### Sensor-space RSA

To track representational organization across electrode space, we additionally repeated the RSA across the six electrode clusters also used in the ERP analysis (see above). Including central electrodes (Fpz, AFz, Fz; FCz, Cz, CPz; Pz, POz, Oz, Iz) in both left- and right-hemispheric clusters yielded electrode counts of 9, 15, and 13, for the anterior, central, and posterior clusters, respectively. All technical details of the cluster-specific RSAs were identical to the analysis using all available electrodes.

#### Controlling for image similarity

To quantify similarity on the image level, we computed pixel similarities for all pairs of images. Each image (220×220 pixels in 3 color layers) was first unfolded into a vector; these vectors were then correlated for each pair of images. A pixel RDM was generated by using 1 – correlation as the dissimilarity measure. As the pixel RDM explained some variance in the face identity RDM (R^2^=.06), neural identity representations could in principle partly reflect pixel similarities. Hence, we used a partial correlation approach (Cichy et al., 2017; Groen et al., 2018), where we repeated the key analyses while removing the pixel RDM by partialing it out. This analysis revealed representations of face identity that are invariant to pixel-based image similarities.

### Statistical testing

To identify significant effects across time, we used a threshold-free cluster enhancement procedure (Smith and Nichols, 2009)with default parameters. Multiple-comparison correction across time was based on a sign-permutation test (with null distributions created from 10,000 bootstrapping iterations) as implemented in CoSMoMVPA (Oosterhof et al., 2016). The resulting statistical maps were thresholded at Z>1.64 (i.e., p<.05, one sided against zero).

## Results

### Event-related potentials reflect face sex, but not face identity

Following traditional EEG studies on face perception, we first performed a univariate ERP analysis across six electrode clusters (Fig. 2). ERPs were different for the four identities primarily in the bilateral posterior electrode clusters (main effect of identity in a four-way ANOVA, Fig. 2E/F, purple line) starting from 100ms for the left and 120ms for the right hemisphere (Fig. 2), remaining significant throughout the length of the epoch. The other electrode clusters showed weaker and less temporally persistent differences (A-D). The difference between identities however originated from the significantly different ERPs for female and male faces from 190ms (left) and 150ms (right), throughout the length of the epoch. By contrast, within-sex comparisons led to no significant results at any of the time-points over either of the electrode clusters. These results support prior studies showing that ERP signals more prominently reflect face sex than face identity (Mouchetant-Rostaing et al., 2000; Freeman et al., 2010). In the following we applied multivariate pattern analysis to further probe the emergence of identity information with higher sensitivity.

**Figure 2.**
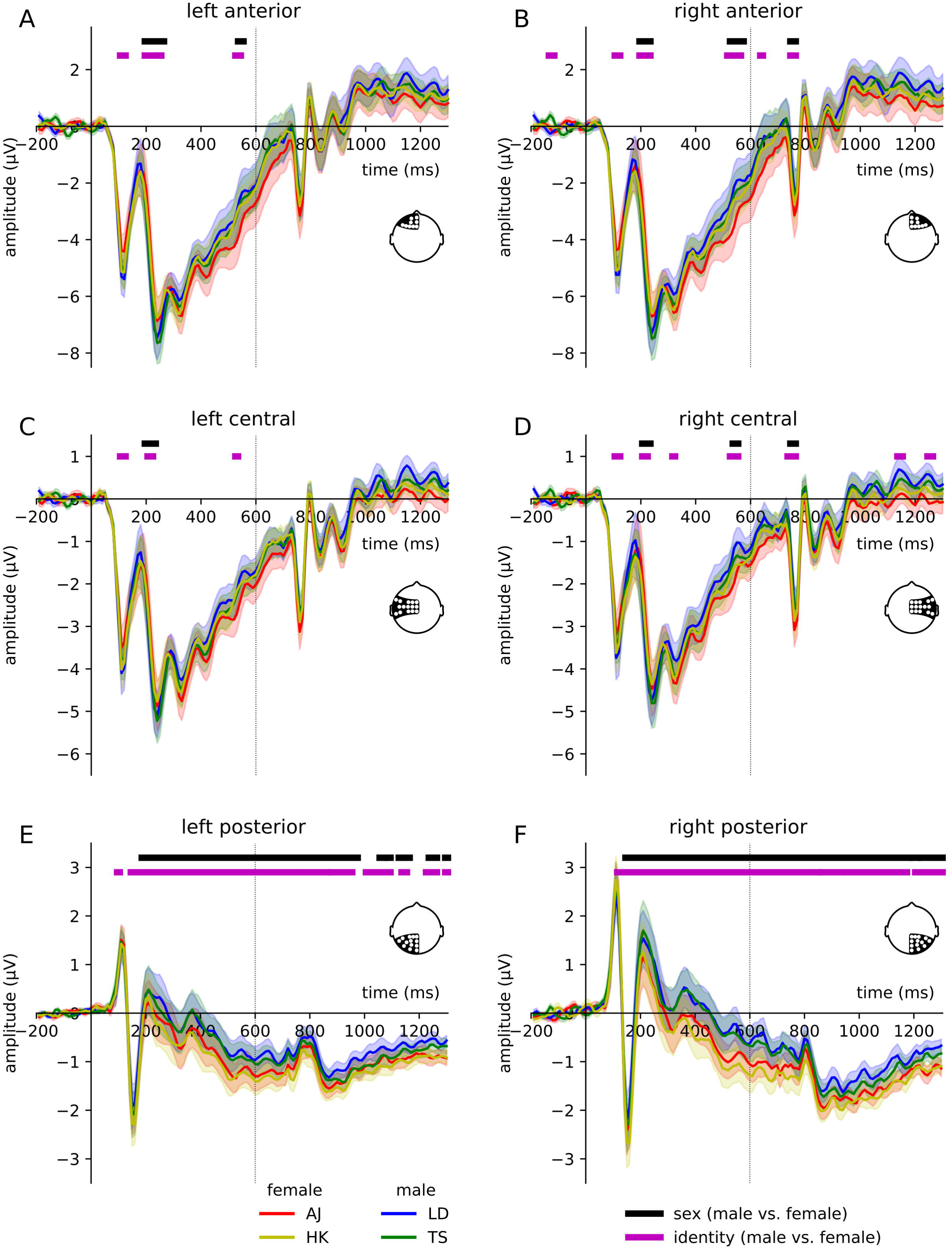
ERP Results. Grand-average ERPs were significantly differed across the four identities (purple significance markers), particularly in the posterior electrode clusters (E, F). This effect was driven by a pronounced difference between male and female faces (black significance markers). Identities of the same sex were not discriminable from ERP responses in any of the clusters, suggesting that ERPs primarily reflect face sex, rather than face identity. Red, blue, yellow and green show the average ERPs for the four celebrities. Horizontal lines denote statistical significance (p<0.05, FDR-corrected for multiple comparisons). Shaded ranges denote standard errors of the mean.

### Tracking the emergence of face identity representations

To reveal identity information in the EEG signals, we generated an identity predictor RDM, which reflected the 40 images’ dissimilarity in identity (Fig. 3A). We then correlated the neural RDM with this identity RDM separately at every time point. This analysis revealed significant correlations from 110ms onwards, peaking at around 410ms (peak t[25]=5.97) and lasting across the whole epoch (Fig. 3B), suggesting rapidly emerging and long-lasting face identity information in the signal.

**Figure. 3:**
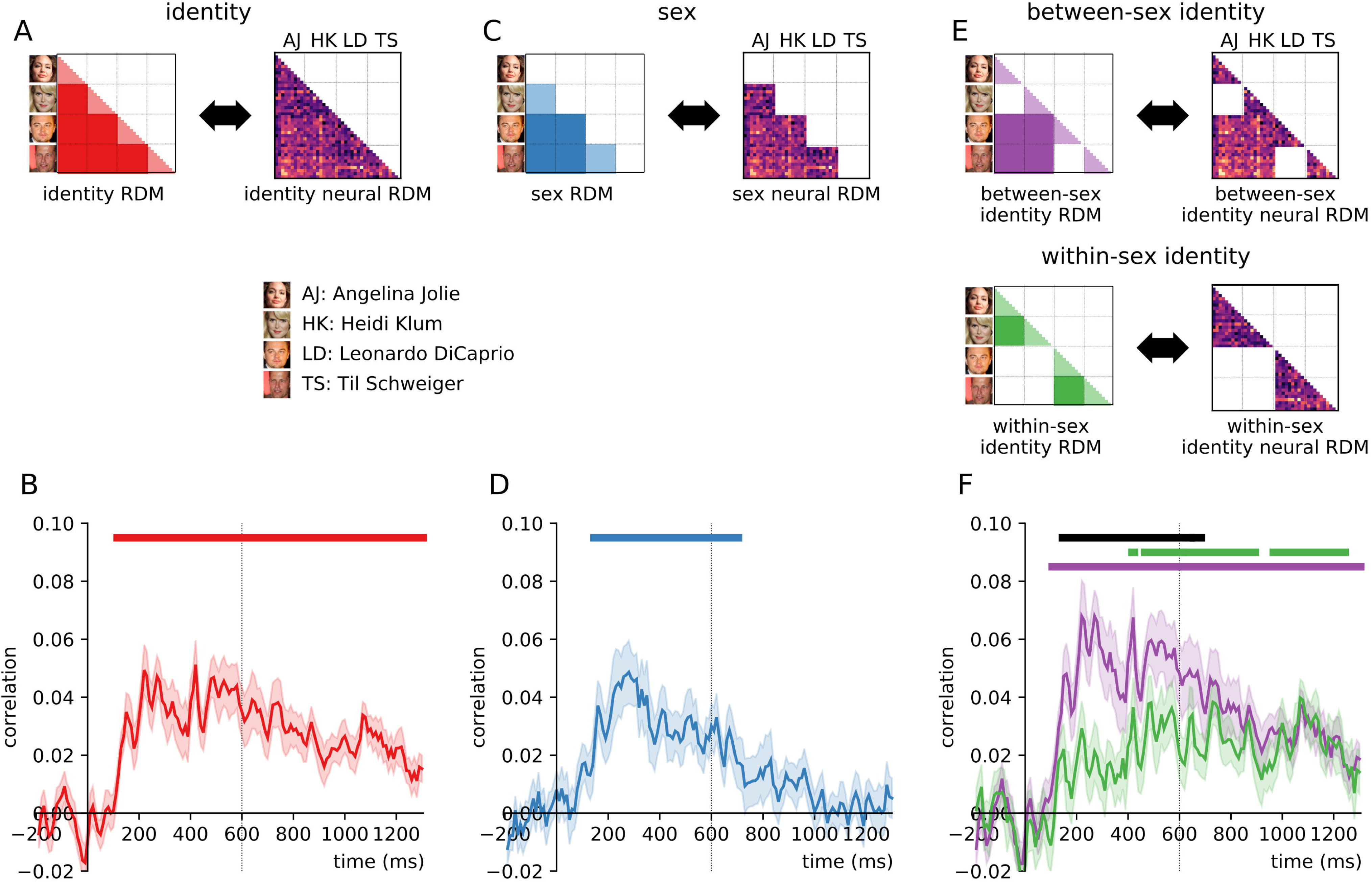
RSA Results. To reveal identity-specific representations, we modeled the representational organization obtained from EEG signals with different predictor matrices (A, C, E). We observed temporally persisting identity information starting from 110ms after stimulus onset (A, B). Similarly, we found strong sex information in the neural organization, emerging between 140ms and 680ms (C, D). Tracking identity information for faces of same and opposite sexes revealed that identity information for same-sex faces was relatively delayed, emerging only after 400ms (E, F). Early identity information was significantly reduced for between-sex comparisons (black significance markers), suggesting that early identity coding partly relies on differences in face sex. Horizontal lines denote statistical significance (p<0.05, corrected for multiple comparisons). Shaded ranges denote standard errors of the mean.

Our stimulus set contained faces of both sexes, and faces within the same sex share more visual and conceptual properties than faces of opposite sexes (O’Toole et al., 1998). To determine whether such sex differences could be retrieved from the EEG signals, we correlated the neural RDM with a sex predictor RDM separately at every time point (Fig. 3C). This sex predictor RDM only contained between-identity comparisons, so that this analysis reflected face sex independently of identity. We found significant sex information from 140ms to 680ms, peaking at 270ms (peak t[25]=4.39) (Fig. 3D). This indicates that the early EEG signals also contain reliable differences between sexes, emerging at a similar time point as identity-specific information but decaying more rapidly.

The presence of sex information in the signal suggests that identity information may be processed differently as a function of the sex of the face. Specifically, as faces of the same sex are more similar in various aspects (including their visual appearance), discriminating between the four facial identities may overestimate the amount of genuine identity information in the signal. We thus split our analysis into comparisons between faces of opposite and of the same sex by correlating the neural RDMs with two separate predictor RDMs (Fig. 3E).

For one of these predictor RDMs (“between-sex”) we only included comparisons between the two sexes, while for the other RDM (“within-sex”) we only included comparisons within the same sex. We observed strong identity information for opposite-sex faces that could be retrieved from as early as 100ms until the end of the epoch and peaking at 260ms (peak t[25]=7.40). Identity information, however, differed when restricting the analysis to within-sex comparisons: it emerged significantly later, at around 400ms, and peaked at 1,050ms (peak t[25]=5.97) (Fig. 3F). When directly comparing identity information for the between- and within-sex comparisons, we found significantly higher identity information for the between-sex analysis between 140ms and 660ms. This suggests that early identity representations partly reflect differences in face sex. By contrast, after 660ms, face sex did not influence identity representations, suggesting the emergence of identity representations that are invariant to commonalities and differences across the two sexes.

### Face identity information predominantly originates from right posterior sources

As highlighted by previous neuroimaging studies (Rossion et al., 2003, 2012) (for a recent review see Yovel, 2016), and evident from our univariate results (see above), face-selective responses are strongest over right posterior electrodes. Using response patterns across the whole scalp may therefore partly obscure face identity information in the multivariate analyses. We thus repeated the RSA separately for each of the six electrode clusters used in the univariate analysis, expecting the strongest identity information in the right posterior cluster (Fig. 4).

**Figure 4.**
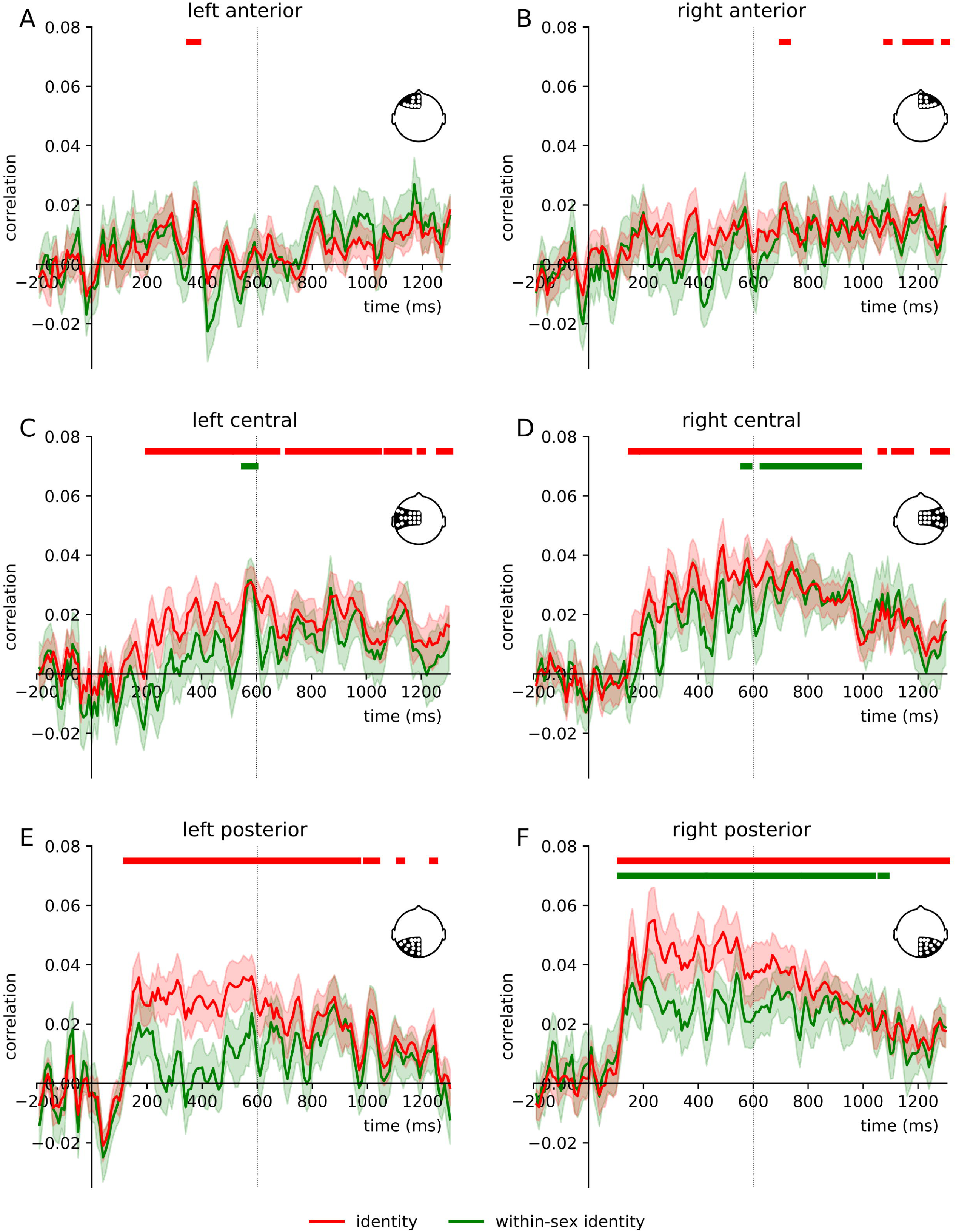
Sensor-space RSA Results. When repeating the RSA for the six electrode clusters used in the ERP analysis, we found strongest identity information in the posterior clusters (E/F). This identity information was lateralized to the right hemisphere: In the right central and posterior electrode clusters (D/F), we observed significant within-sex identity information, with an early onset (110ms) in the right posterior cluster. The corresponding left-hemispheric clusters (C/E) only yielded identity information when also the between-sex comparisons were included. The anterior clusters (A/B) did not yield substantial identity information. Horizontal lines denote statistical significance (p<0.05, corrected for multiple comparisons). Shaded ranges denote standard errors of the mean.

For the posterior electrode clusters we found the most pronounced identity information, and a marked difference between hemispheres. In the left posterior cluster, four-way identity information (where sex may contribute to identity encoding) emerged from 120ms post-stimulus onset and peaked at 560ms (peak t[25]=4.85) (Fig. 4E). However, restricting the analysis to within-sex comparisons abolished identity information over this electrode cluster in the signal entirely. Similarly, in the right posterior cluster (Fig. 4F) we found robust four-way identity information, starting from 110ms after stimulus onset and peaking at 230ms (peak t[25]=4.81). Crucially however, the right posterior cluster also showed reliable within-sex identity information throughout the epoch, emerging at the same time, after 110ms and peaking around 530ms (peak t[25]=5.40). This result suggests that signals recorded from electrodes close to the typically face-selective ERP recording sites of the right hemisphere contain widespread identity information, even when visual and conceptual properties are more robustly controlled for.

The left central cluster (Fig. 4C) primarily showed four-way identity information, emerging slightly later as compared to the posterior cluster, after 200ms, peaking at 560ms (peak t[25]=5.10). By contrast, the right central cluster not only yielded four-way identity information (from 150ms, peaking at 480ms, peak t[25]=4.97), but also within-sex identity information, emerging later than that of the posterior cluster, after 550ms and peaking at 1,100ms (peak t[25]=3.11).

Signals recorded from the two anterior clusters did not yield substantial identity information (Fig. 4A/B), suggesting that identity information primarily originates from sources in visual cortex.

### Late representations of face identity are invariant to image properties

Our stimulus set was constructed to mirror natural variations across different encounters with a familiar person. This was achieved by selecting stimuli that ensured a high degree of variability within each identity (see above), so that image-based stimulus properties are unlikely to account for the emergence of identity information. To explicitly rule out this possibility, we performed a control analysis, where we additionally modeled image-based similarities between stimuli. This was done by constructing pixel RDMs, which reflected the images’ dissimilarity in pixel values; these pixel RDMs were partialled out in the subsequent analysis. We focused the control analysis on the within-sex comparison, which forms the most robust test of face identity representations, and on the two electrode configurations where it was most robustly found (all electrodes and right posterior electrodes).

In the analysis using all electrodes, we found no modulation of identity information after removing the pixel RDM (Fig. 5B). By contrast, when focusing on the right posterior cluster, we found a modulation of identity information when controlling for image-based similarity (Fig. 5C). Early within-sex identity information, emerging between 90ms and 230ms was significantly reduced when controlling for pixel dissimilarity. By contrast, later within-sex identity information (from 460ms) emerged independently of image-based properties. Together, these results suggest that later representations of face identity are robust to image-based changes, but genuinely reflect face identity. These neural representations might thus be a crucial prerequisite for efficient face recognition across visually different encounters with a person.

**Figure 5.**
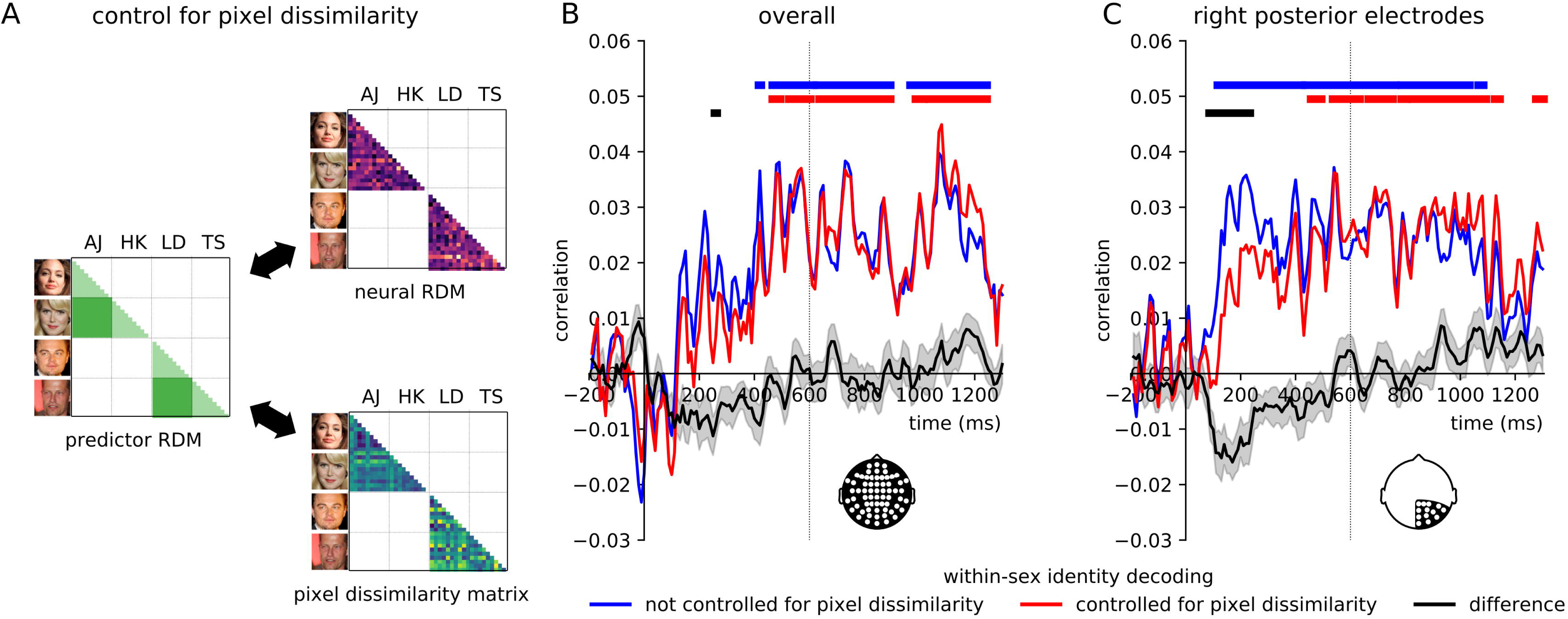
Controlling for Image Similarity. In a partial correlation analysis, we tracked within-sex identity information while controlling for the images’ pixel dissimilarities (A). When using data from all electrodes, removing pixel dissimilarities did not significantly impact identity information (B). For the right posterior cluster, where early within-sex identity information was found in previous analysis (Fig. 4F), controlling for pixel dissimilarity had a significant impact (C): early identity information (90-230ms) was significantly reduced when controlling for pixel dissimilarity, whereas later identity information was not impacted and remained significant from 460ms after onset. These results suggest that later representations of face identity are invariant to image-based properties. Horizontal lines denote statistical significance (p<0.05, corrected for multiple comparisons). Shaded ranges denote standard errors.

## Discussion

In the current study, we applied representational similarity analysis to EEG signals to investigate the neural dynamics of familiar face recognition. Our results show that face identity can be rapidly recovered from EEG response patterns, even with highly variable, “ambient” face stimuli (Jenkins et al., 2011). In more fine-grained analyses, we uncovered a gradual emergence of face identity coding: Early identity information is modulated by face sex and by visual image properties. By contrast, later identity information, emerging after 400ms and primarily in the right hemisphere, is unaffected by these factors. This finding suggests that after 400ms representations genuinely reflect face identity. These later representations may be the basis for real-world face recognition, allowing the identification of an individual across different encounters and against similar-looking other faces.

In everyday life, the facial appearance of a single person can be highly variable. This variability makes it challenging to match an individual encounter with a face to an identity representation stored in memory (Bruce et al., 1999; Clutterbuck and Johnston, 2002; Jenkins et al., 2011; Andrews et al., 2015). The invariant identity representations revealed here are ideal for extracting face identity from different encounters, as they discriminate identities of the same sex, across variations in visual properties. The late emergence of such representations is compatible with the involvement of conceptual identity representations in the medial and anterior temporal cortices (Quiroga et al., 2005; Mormann et al., 2008); linking our EEG results with functional neuroimaging data (Cichy et al., 2014, 2016) could directly test this possibility in the future.

How do these seemingly late identity representations support rapid face recognition in the wild? While these representations are useful under great variability and in the presence of distracting face information, face recognition is sometimes easier than this: In real-life situations, we often know which person to expect, which visual properties are diagnostic of him or her, and where the person likely shows up. Under such conditions, motor responses in face recognition tasks can be faster than 400ms (Besson et al., 2017). This observation suggests that face identity can sometimes be inferred from earlier representations that do not need to be highly invariant. Future studies could thus test whether different representational stages are crucial for face recognition under varying demands.

Our study revealed a pronounced right-hemispheric lateralization of identity information: face identity information was strongest in electrodes over the right, as opposed to the left, visual cortex. Specifically, only signals recorded over right occipito-temporal cortex contained identity information which is invariant to both face sex and image-based properties. This right-lateralized topography is consistent with sources in the visual face processing network that has a strong right-hemispheric lateralization (Axelrod and Yovel, 2015; Yovel, 2016). Interestingly, neuroimaging work showed that specifically right-hemispheric activations predict behavioral performance in familiar face recognition (Weibert and Andrews, 2015), suggesting that these identity representations could play an important role in face recognition. However, this notion has to be explicitly tested in the future, as caution needs to be applied when inferring cortical sources from EEG scalp topographies.

Besides identity coding, our findings also offer insights into the cortical coding of face sex. As our stimulus set contained faces of opposite sexes, we could also track the emergence of sex information. Face sex can be rapidly retrieved from EEG signals, both in univariate and multivariate analyses, and predicts cortical organization from 140ms. This finding corroborates previous ERP studies, which have suggested that face sex is extracted early and affects a variety of face-related ERP components (Mouchetant-Rostaing et al., 2000; Ito and Urland, 2003, 2005; Kloth et al., 2015). As opposed to the temporally sustained identity information, sex information displayed a more transient nature, and vanished shortly before 700ms after onset. This difference between identity and sex information suggests that the two properties are coded somewhat independently at later processing stages.

In conclusion, we provide a characterization of the neural dynamics underlying familiar face recognition. Representations of face identity emerged gradually across the visual processing cascade. Invariant identity representations were observed after 400ms of processing. We suggest that these representations are crucial for face recognition across different encounters with a person.

## Footnotes

1 Image credits: <u>File:Angelina</u> Jolie at Davos crop.jpg. (2014, April 23). *Wikimedia Commons, the free media repository*. Retrieved 15:17, May 1, 2018 from https://commons.wikimedia.org/w/index.php?title=File:Angelina_Jolie_at_Davos_crop.jpg&oldid=122076100. Creative Commons Attribution-Share Alike 3.0 Unported license. <u>File:LeonardoDiCaprioNov08.jpg</u>. (2018, January 20). *Wikimedia Commons, the free media repository*. Retrieved 15:18, May 1, 2018 from https://commons.wikimedia.org/w/index.php?title=File:LeonardoDiCaprioNov08.jpg&oldid=281183411. Creative Commons Attribution-Share Alike 3.0 Unported license. These images were not part of the original stimulus set.

## Acknowledgements

The authors would like to thank Lisa-Celine Süllwold and Bettina Kamchen for their help in participant recruitment and data acquisition. This work was supported by grants from the Deutsche Forschungsgemeinschaft (KO3918/5-1, KA4683-2/1, CI241-1/1).

## References

Andrews S, Jenkins R, Cursiter H, Burton AM, Andrews S, Jenkins R, Cursiter H, Telling AMB, Andrews S, Jenkins R, Cursiter H, Burton AM (2015) Telling faces together : Learning new faces through exposure to multiple instances Telling faces together : Learning new faces through exposure to multiple instances. Q J Exp Psychol 0218:2041–2050 Available at: https://www.ncbi.nlm.nih.gov/pubmed/25607814.

Anzellotti S, Caramazza A (2014) The neural mechanisms for the recognition of face identity in humans. Front Psychol 5:672 Available at: https://www.ncbi.nlm.nih.gov/pmc/articles/PMC4072087/.

Anzellotti S, Fairhall SL, Caramazza A (2014) Decoding representations of face identity that are tolerant to rotation. Cereb Cortex 24:1988–1995 Available at: https://academic.oup.com/cercor/article-lookup/doi/10.1093/cercor/bht046.

Axelrod V, Yovel G (2015) Successful decoding of famous faces in the fusiform face area. PLoS One 10:e0117126 Available at: http://journals.plos.org/plosone/article?id=10.1371/journal.pone.0117126.

Bentin S, Deouell LY (2000) Structural Encoding and Identification in Face Processing: Erp Evidence for Separate Mechanisms. Cogn Neuropsychol 17:35–55 Available at: http://www.tandfonline.com/doi/abs/10.1080/026432900380472.

Besson G, Barragan-Jason G, Thorpe SJ, Fabre-Thorpe M, Puma S, Ceccaldi M, Barbeau EJ (2017) From face processing to face recognition: Comparing three different processing levels. Cognition 158:33–43 Available at: http://www.sciencedirect.com/science/article/pii/S0010027716302384.

Bruce V, Henderson Z, Greenwood K, Hancock PJB, Burton AM, Miller P (1999) Verification of face identities from images captured on video. J Exp Psychol Appl 5:339–360 Available at: http://psycnet.apa.org/fulltext/1999-01801-001.html.

Caharel S, Jiang F, Blanz V, Rossion B (2009) Recognizing an individual face: 3D shape contributes earlier than 2D surface reflectance information. Neuroimage 47:1809–1818 Available at: https://www.ncbi.nlm.nih.gov/pubmed/19497375.

Cichy RM, Kriegeskorte N, Jozwik KM, Bosch JJF van den, Charest I (2017) Neural dynamics of real-world object vision that guide behaviour. bioRxiv:147298 Available at: https://www.biorxiv.org/content/early/2017/06/08/147298.

Cichy RM, Pantazis D, Oliva A (2014) Resolving human object recognition in space and time. Nat Neurosci 17:455–462 Available at: https://www.nature.com/articles/nn.3635.

Cichy RM, Pantazis D, Oliva A (2016) Similarity-Based Fusion of MEG and fMRI Reveals Spatio-Temporal Dynamics in Human Cortex During Visual Object Recognition. Cereb Cortex 26:3563–3579 Available at: https://academic.oup.com/cercor/article/26/8/3563/2428700.

Clutterbuck R, Johnston RA (2002) Exploring levels of face familiarity by using an indirect face-matching measure. Perception 31:985–994 Available at: https://www.ncbi.nlm.nih.gov/pubmed/12269591.

Curran T, Hancock J (2007) The FN400 indexes familiarity-based recognition of faces. Neuroimage 36:464–471 Available at: https://www.sciencedirect.com/science/article/pii/S1053811906011839.

Debruille JB, Guillem F, Renault B (1998) ERPs and chronometry of face recognition: following-up Seeck et al. and George et al. Neuroreport 9:3349–3353 Available at: http://www.ncbi.nlm.nih.gov/pubmed/9855278 [Accessed July 7, 2018].

Duchaine B, Yovel G (2015) A Revised Neural Framework for Face Processing. Annu Rev Vis Sci 1:393–416 Available at: http://www.annualreviews.org/doi/10.1146/annurev-vision-082114-035518.

Freeman JB, Ambady N, Holcomb PJ (2010) The face-sensitive N170 encodes social category information. Neuroreport Available at: https://www.ncbi.nlm.nih.gov/pmc/articles/PMC3576572.

Gilaie-Dotan S, Malach R (2007) Sub-exemplar shape tuning in human face-related areas. Cereb Cortex 17:325–338 Available at: https://academic.oup.com/cercor/article-pdf/17/2/325/17297436/bhj150.pdf.

Gobbini MI, Haxby J V. (2007) Neural systems for recognition of familiar faces. Neuropsychologia 45:32– Available at: https://www.ncbi.nlm.nih.gov/pubmed/16797608.

Goesaert E, Op de Beeck HP (2013) Representations of Facial Identity Information in the Ventral Visual Stream Investigated with Multivoxel Pattern Analyses. J Neurosci 33:8549–8558 Available at: http://www.jneurosci.org/cgi/doi/10.1523/JNEUROSCI.1829-12.2013.

Gosling A, Eimer M (2011) An event-related brain potential study of explicit face recognition. Neuropsychologia 49:2736–2745.

Gramfort A, Luessi M, Larson E, Engemann DA, Strohmeier D, Brodbeck C, Goj R, Jas M, Brooks T, Parkkonen L, Hämäläinen M (2013) MEG and EEG data analysis with MNE-Python. Front Neurosci Available at: https://www.frontiersin.org/articles/10.3389/fnins.2013.00267/full.

Gramfort A, Luessi M, Larson E, Engemann DA, Strohmeier D, Brodbeck C, Parkkonen L, Hämäläinen MS (2014) MNE software for processing MEG and EEG data. Neuroimage 86:446–460 Available at: https://www.ncbi.nlm.nih.gov/pubmed/24161808.

Groen IIA, Greene MR, Baldassano C, Fei-Fei L, Beck DM, Baker CI (2018) Distinct contributions of functional and deep neural network features to representational similarity of scenes in human brain and behavior. Elife 7 Available at: https://elifesciences.org/articles/32962.

Grootswagers T, Wardle SG, Carlson TA (2017) Decoding dynamic brain patterns from evoked responses: A tutorial on multivariate pattern analysis applied to time series neuroimaging data. J Cogn Neurosci 29:677–697 Available at: https://www.mitpressjournals.org/doi/abs/10.1162/jocn_a_01068.

Heisz JJ, Watter S, Shedden JM (2006) Automatic face identity encoding at the N170. Vision Res 46:4604–4614 Available at: https://www.sciencedirect.com/science/article/pii/S0042698906004573.

Huddy V, Schweinberger SR, Jentzsch I, Burton AM (2003) Matching faces for semantic information and names: an event-related brain potentials study. Cogn Brain Res 17:314–326 Available at: https://www.sciencedirect.com/science/article/pii/S0926641003001319 [Accessed July 7, 2018].

Ito TA, Urland GR (2003) Race and Gender on the Brain: Electrocortical Measures of Attention to the Race and Gender of Multiply Categorizable Individuals. J Pers Soc Psychol 85:616–626 Available at: https://www.ncbi.nlm.nih.gov/pubmed/14561116.

Ito TA, Urland GR (2005) The influence of processing objectives on the perception of faces: An ERP study of race and gender perception. Cogn Affect Behav Neurosci 5:21–36 Available at: https://www.ncbi.nlm.nih.gov/pubmed/15913005.

Jas M, Engemann DA, Bekhti Y, Raimondo F, Gramfort A (2017) Autoreject: Automated artifact rejection for MEG and EEG data. Neuroimage 159:417–429 Available at: https://doi.org/10.1016/j.neuroimage.2017.06.030.

Jenkins R, White D, Van Montfort X, Mike Burton A (2011) Variability in photos of the same face. Cognition 121:313–323 Available at: https://www.ncbi.nlm.nih.gov/pubmed/21890124.

Jin J, Allison BZ, Kaufmann T, Kübler A, Zhang Y, Wang X, Cichocki A (2012) The Changing Face of P300 BCIs: A Comparison of Stimulus Changes in a P300 BCI Involving Faces, Emotion, and Movement Frishman L, ed. PLoS One 7:e49688 Available at: http://dx.plos.org/10.1371/journal.pone.0049688 [Accessed July 7, 2018].

Johnston R a, Edmonds AJ (2009) Familiar and unfamiliar face recognition: a review. Memory 17:577–596 Available at: http://www.ncbi.nlm.nih.gov/pubmed/19548173.

Kaiser D, Oosterhof NN, Peelen M V. (2016) The Neural Dynamics of Attentional Selection in Natural Scenes. J Neurosci 36:10522–10528 Available at: http://www.jneurosci.org/cgi/doi/10.1523/JNEUROSCI.1385-16.2016.

Kloth N, Damm M, Schweinberger SR, Wiese H (2015) Aging affects sex categorization of male and female faces in opposite ways. Acta Psychol (Amst) 158:78–86 Available at: https://www.sciencedirect.com/science/article/pii/S0001691815000931 [Accessed July 7, 2018].

Kramer RSS, Young AW, Burton AM (2018) Understanding face familiarity. Cognition 172:46–58 Available at: https://www.ncbi.nlm.nih.gov/pubmed/29232594.

Kriegeskorte N (2008) Representational similarity analysis – connecting the branches of systems neuroscience. Front Syst Neurosci 2:4 Available at: http://journal.frontiersin.org/article/10.3389/neuro.06.004.2008/abstract [Accessed July 7, 2018].

Kriegeskorte N, Formisano E, Sorger B, Goebel R (2007) Individual faces elicit distinct response patterns in human anterior temporal cortex. Proc Natl Acad Sci 104:20600–20605 Available at: http://www.pnas.org/cgi/doi/10.1073/pnas.0705654104.

Kriegeskorte N, Kievit RA (2013) Representational geometry: Integrating cognition, computation, and the brain. Trends Cogn Sci 17:401–412 Available at: https://www.ncbi.nlm.nih.gov/pmc/articles/PMC3730178/.

Liu J, Harris A, Kanwisher N (2013) Stages of processing in face perception: An MEG study. Soc Neurosci Key Readings 5:75–86 Available at: http://www.nature.com/articles/nn909 [Accessed July 7, 2018].

Mike Burton A (2013) Why has research in face recognition progressed so slowly? The importance of variability. Q J Exp Psychol 66:1467–1485 Available at: http://www.ncbi.nlm.nih.gov/pubmed/23742022.

Mormann F, Kornblith S, Quiroga RQ, Kraskov A, Cerf M, Fried I, Koch C (2008) Latency and Selectivity of Single Neurons Indicate Hierarchical Processing in the Human Medial Temporal Lobe. J Neurosci 28:8865–8872 Available at: http://www.jneurosci.org/cgi/doi/10.1523/JNEUROSCI.1640-08.2008.

Mouchetant-Rostaing Y, Giard MH, Bentin S, Aguera PE, Pernier J (2000) Neurophysiological correlates of face gender processing in humans. Eur J Neurosci.

Nasr S, Tootell RBH (2012) Role of fusiform and anterior temporal cortical areas in facial recognition. Neuroimage 63:1743–1753 Available at: https://www.ncbi.nlm.nih.gov/pubmed/23034518.

Natu V, O’Toole AJ (2011) The neural processing of familiar and unfamiliar faces: A review and synopsis. Wiley/Blackwell (10.1111). Available at: http://doi.wiley.com/10.1111/j.2044-8295.2011.02053.x [Accessed July 7, 2018].

Nemrodov D, Niemeier M, Mok JNY, Nestor A (2016) The time course of individual face recognition: A pattern analysis of ERP signals. Neuroimage 132:469–476 Available at: http://linkinghub.elsevier.com/retrieve/pii/S1053811916002020 [Accessed July 7, 2018].

Nemrodov D, Niemeier M, Patel A, Nestor A (2018) The Neural Dynamics of Facial Identity Processing: insights from EEG-Based Pattern Analysis and Image Reconstruction. eneuro:ENEURO.0358-17.2018 Available at: http://eneuro.sfn.org/lookup/doi/10.1523/ENEURO.0358-17.2018 [Accessed July 7, 2018].

Nestor A, Plaut DC, Behrmann M (2011) Unraveling the distributed neural code of facial identity through spatiotemporal pattern analysis. Proc Natl Acad Sci 108:9998–10003 Available at: http://www.pnas.org/cgi/doi/10.1073/pnas.1102433108.

O’Toole AJ, Deffenbacher KA, Valentin D, McKee K, Huff D, Abdi H (1998) The perception of face gender: The role of stimulus structure in recognition and classification. Mem Cogn 26:146–160 Available at: https://www.ncbi.nlm.nih.gov/pubmed/9519705.

Oosterhof NN, Connolly AC, Haxby J V. (2016) CoSMoMVPA: Multi-Modal Multivariate Pattern Analysis of Neuroimaging Data in Matlab/GNU Octave. Front Neuroinform 10:27 Available at: http://journal.frontiersin.org/Article/10.3389/fninf.2016.00027/abstract [Accessed July 7, 2018].

Peirce JW (2008) Generating stimuli for neuroscience using PsychoPy. Front Neuroinform 2 Available at: http://journal.frontiersin.org/article/10.3389/neuro.11.010.2008/abstract.

Quiroga RQ, Reddy L, Kreiman G, Koch C, Fried I (2005) Invariant visual representation by single neurons in the human brain. Nature 435:1102–1107 Available at: https://www.nature.com/articles/nature03687.

Rossion B, Hanseeuw B, Dricot L (2012) Defining face perception areas in the human brain: A large-scale factorial fMRI face localizer analysis. Brain Cogn 79:138–157 Available at: https://www.ncbi.nlm.nih.gov/pubmed/22330606.

Rossion B, Jacques C (2008) Does physical interstimulus variance account for early electrophysiological face sensitive responses in the human brain? Ten lessons on the N170. Neuroimage 39:1959–1979 Available at: https://www.ncbi.nlm.nih.gov/pubmed/18055223.

Rossion B, Joyce CA, Cottrell GW, Tarr MJ (2003) Early lateralization and orientation tuning for face, word, and object processing in the visual cortex. Neuroimage 20:1609–1624 Available at: https://www.ncbi.nlm.nih.gov/pubmed/14642472.

Rousselet GA, Husk JS, Pernet CR, Gaspar CM, Bennett PJ, Sekuler AB (2009) Age-related delay in information accrual for faces: Evidence from a parametric, single-trial EEG approach. BMC Neurosci 10:114 Available at: http://bmcneurosci.biomedcentral.com/articles/10.1186/1471-2202-10-114 [Accessed July 7, 2018].

Schweinberger SR, Pickering EC, Jentzsch I, Burton AM, Kaufmann JM (2002) Event-related brain potential evidence for a response of inferior temporal cortex to familiar face repetitions. Cogn Brain Res 14:398–409 Available at: https://www.ncbi.nlm.nih.gov/pubmed/12421663.

Smith SM, Nichols TE (2009) Threshold-free cluster enhancement: Addressing problems of smoothing, threshold dependence and localisation in cluster inference. Neuroimage 44:83–98 Available at: https://www.ncbi.nlm.nih.gov/pubmed/18501637.

Tanaka JW, Curran T, Porterfield AL, Collins D (2006) Activation of Preexisting and Acquired Face Representations: The N250 Event-related Potential as an Index of Face Familiarity. J Cogn Neurosci 18:1488–1497 Available at: http://www.mitpressjournals.org/doi/10.1162/jocn.2006.18.9.1488 [Accessed July 7, 2018].

Verosky SC, Todorov A, Turk-Browne NB (2013) Representations of individuals in ventral temporal cortex defined by faces and biographies. Neuropsychologia 51:2100–2108 Available at: https://www.ncbi.nlm.nih.gov/pubmed/23871881.

Vida MD, Nestor A, Plaut DC, Behrmann M (2017) Spatiotemporal dynamics of similarity-based neural representations of facial identity. Proc Natl Acad Sci 114:388–393 Available at: http://www.pnas.org/lookup/doi/10.1073/pnas.1614763114 [Accessed July 7, 2018].

Visconti Di Oleggio Castello M, Halchenko YO, Guntupalli JS, Gors JD, Gobbini MI (2017) The neural representation of personally familiar and unfamiliar faces in the distributed system for face perception. Sci Rep 7:12237 Available at: http://www.nature.com/articles/s41598-017-12559-1 [Accessed July 7, 2018].

Wang Z, Bovik AC, Sheikh HR, Simoncelli EP (2004) Image quality assessment: From error visibility to structural similarity. IEEE Trans Image Process 13:600–612 Available at: https://ieeexplore.ieee.org/document/1284395.

Weibert K, Andrews TJ (2015) Activity in the right fusiform face area predicts the behavioural advantage for the perception of familiar faces. Neuropsychologia 75:588–596 Available at: https://www.ncbi.nlm.nih.gov/pubmed/26187507.

Weibert K, Harris RJ, Mitchell A, Byrne H, Young AW, Andrews TJ (2016) An image-invariant neural response to familiar faces in the human medial temporal lobe. Cortex 84:34–42 Available at: https://www.ncbi.nlm.nih.gov/pubmed/27697662.

Young AW, Burton AM (2017) Recognizing Faces. Curr Dir Psychol Sci 26:212–217 Available at: journals.sagepub.com/doi/abs/10.1177/0963721416688114.

Yovel G (2016) Neural and cognitive face-selective markers: An integrative review. Neuropsychologia 83:5–13 Available at: https://www.ncbi.nlm.nih.gov/pubmed/26407899.

